## Supplementary information for "An interbacterial lipase toxin with an unprecedented reverse domain arrangement defines a new class of type VII secretion system effector"

### Supplementary Material

| | USA300 positive control | USA300 negative control | USA300 TslA-TlaA1-TlaA2 | USA300 Δ*til1* positive control | USA300 Δ*til1* negative control | USA300 Δ*til1* TslA-TlaA1-TlaA2 |
|---|---|---|---|---|---|---|
| **Total cells** | 1216 | 680 | 1538 | 1346 | 735 | 864 |
| **Total cells with membrane damage** | 560 | 8 | 3 | 905 | 1 | 179 |
| **Percentage cells membrane damage** | 46.05 | 1.18 | 0.20 | 67.24 | 0.14 | 20.72 |

**Supplementary Table 1. Analysis of *S. aureus* USA300 cells stained by Sytox green when imaged using fluorescence microscopy.** The percentage cells stained with Sytox green was calculated for each strain from the total cells analysed, as described in the methods. Data is represented in Fig. 5e.

| Strain | Relevant genotype or description | Source or reference |
|---|---|---|
| *E. coli* | | |
| JM110 | *rpsL thr leu thi lacY galK galT ara tonA tsx dam dcm glnV44 Δ(lac-proAB) e14-* [*F' traD36 proAB+ lacIq lacZΔM15*] *hsdR17(rK−mK+)* | Stratagene |
| TOP10 | *F- mcrA (mrr-hsdRMS-mcrBC) 80lacZ M15 lacX74 recA1 ara 139 (ara-leu)7697 galU galK rpsL (*Str$^{r}$*) endA1 nupG* | Invitrogen |
| BL21(DE3) | *E. coli B: F$^{-}$, dcm, ompT, hsdS(rB-, mB-), gal,* λ(DE3) | Reference[1] |
| M15 [pREP4] | *F-, lac, ara, gal, mtl,* [Kan$^{r}$, *lacI*] | Qiagen |
| BTH101 | F$^{-}$ *cya-99, araD139, galE15, galK16, rpsL1* (Str$^{r}$), *hsdR2, mcrA1, mcrB1* | Reference[2] |
| DH5a | φ80d *Δ(lacZ)M15 recA1 endA1 gyrA96 thi-1 hsdR17 (rk-mk+) supE44 relA1 deoR* Δ(*lacZYA-argF)*U169 | Promega |
| *S. aureus* | | |
| RN6390 | NCTC8325 derivative, *rbsU, tcaR*, cured of φ11, φ12, φ13 | Reference[3] |
| USA300 LAC | Wild type | Reference[4] |
| USA300 Δ*essC* | In-frame deletion of *essC* from USA300 LAC | This work |
| USA300 Δ*tslA* | In-frame deletion of *tslA* from USA300 LAC | This work |
| USA300 Δ*til1* | Also named *S. aureus* USA300 Δ*lpl*Δ*lpp3*Δ*lpp4*Δ*csa1*. In-frame deletions of all four LPL loci. | Reference[5] |
| USA300 Δ*til1* Δ*essC* | In-frame deletion of *essC* from USA300 Δ*til1* | This work |
| USA300 Δ*til1::tilA* | As USA300 *Δtil1*, with a copy of *tilA* introduced between *SAOUHSC_00037* and *SAOUHSC_00039* using pTH100_tilA. | This Work |

**Supplementary Table 2.** Bacterial strains used in this work.

| Plasmid | Relevant genotype or description | Source or reference |
|---|---|---|
| pIMAY | *E. coli/S. aureus* shuttle vector, temperature sensitive, cml$^r$ | Reference[6] |
| pIMAY-essC | pIMAY carrying *essC* deletion allele | Reference[7] |
| pIMAY-Z | *E. coli/S. aureus* shuttle vector, temperature sensitive, cml$^r$ | Reference[8] |
| pIMAY-Z-tslA | pIMAY carrying *tslA* deletion allele | This work |
| pTH100 | Plasmid for markerless integration of GFP into *S. aureus* | Reference[9] |
| pTH100-tilA | As pTH100 but *gfp* replaced by *tilA,* including its native ribosome binding site (rbs) | This work |
| pRAB11 | *E. coli/S. aureus* shuttle vector, inducible protein expression, *amp$^r$, cml$^r$* | Reference[10] |
| pRAB11-TslA | pRAB11 encoding *tslA*, preceded by *hla* rbs | This work |
| pRAB11-TslA-TlaA1-TlaA2 | pRAB11 encoding *tslA-tlaA1-tlaA2*. The *hla* rbs precedes *tslA* | This work |
| pRAB11-TslA-TlaA1 | As pRAB11 encoding *tslA-tlaA1-tlaA2*, with the *tlaA2* gene deleted. The *hla* rbs precedes *tslA*. Synthesised by GenScript. | This work |
| pRAB11-TslA -TlaA2 | As pRAB11 encoding *tslA-tlaA1-tlaA2*, with the *tlaA1* gene deleted. The *hla* rbs precedes *tslA.* Synthesised by GenScript. | This work |
| pRAB11-TslA$_{S164A}$-TlaA1-TlaA2 | As pRAB11-TslA-TlaA1-TlaA2 but encoding TslA S164A substitution | This work |
| pRAB11-TslA$_{D224A}$-TlaA1-TlaA2 | As pRAB11-TslA-TlaA1-TlaA2 but encoding TslA D224A substitution | This work |
| pRAB11-TslA$_{H251A}$-TlaA1-TlaA2 | As pRAB11-TslA-TlaA1-TlaA2 but encoding TslA H251A substitution. | This work |
| pRAB11-TslA$_{L312A}$-TlaA1-TlaA2 | As pRAB11-TslA-TlaA1-TlaA2 but encoding TslA L312A substitution. Synthesised by GenScript. | This work |
| pRAB11-TslA$_{G310A}$-TlaA1-TlaA2 | As pRAB11-TslA-TlaA1-TlaA2 but encoding TslA G310A substitution. Synthesised by GenScript. | This work |
| pRAB11-TslA$_{G310S}$-TlaA1-TlaA2 | As pRAB11-TslA-TlaA1-TlaA2 but encoding TslA G310S substitution. Synthesised by GenScript. | This work |
| pRAB11-pep86_EsxA | pRAB11 producing EsxA fused to an N-terminal pep86 tag. The *hla* rbs precedes *esxA* | This Work |
| pRAB11-pep86_TrxA | pRAB11 producing TrxA fused to an N-terminal pep86 tag. The *hla* rbs precedes *trxA* | Reference[11] |
| pRAB11-pep86_TslA | pRAB11 producing TslA fused to an N-terminal pep86 tag. The *hla* rbs precedes *tslA*. Synthesised by GenScript. | This Work |
| pRAB11-pep86_TslA-TlaA1-TlaA2 | pRAB11 encoding *tslA-tlaA1-tlaA2*, producing TslA fused to an N-terminal pep86 tag. The *hla* rbs precedes *tslA* | This Work |
| pRAB11-TslA-TlaA1-pep86_TlaA2 | pRAB11 encoding *tslA-tlaA1-tlaA2*, producing TlaA2 fused to an N-terminal pep86 tag. The *hla* ribosome binding site precedes both *tslA* and *tlaA2* | This Work |
| pRAB11-TslA-TlaA1_pep86-TlaA2 | pRAB11 encoding *tslA-tlaA1-tlaA2*, producing TlaA1 fused to an C-terminal pep86 tag. The *hla* rbs precedes *tslA*. Synthesised by GenScript. | This work |
| pBAD-$_{6H}$11S | Expression vector for purification of 11S | Reference[12] |

| | | |
|---|---|---|
| pQE70 | Vector for regulatable protein overproduction in *E. coli* (T5 promoter). amp$^{r}$ | Qiagen |
| pQE70-TslA-His | pQE70 with *tslA* cloned in frame in the multiple cloning site to produce a C-terminal His(6)-tag fusion. | This Work |
| pQE70-TslA$_{S164A}$-His | As pQE70-TslA-His but encoding TslA S164A substitution | This Work |
| pQE70-TslA$_{D224A}$-His | As pQE70-TslA-His but encoding TslA D224A substitution | This Work |
| pQE70-TslA$_{H251A}$-His | As pQE70-TslA-His but encoding TslA H521A substitution | This Work |
| pREP4 | *lacI* kan$^{r}$ | Qiagen |
| pLysS | Encodes T7 lysozyme cml$^{r}$ | Promega |
| pET15bTEV | Overexpression plasmid, T7 promoter. Adds (His)6-tag and a TEV site to N-terminus of protein | Reference[13] |
| pET15bTEV-TslA$_{CT}$-TlaA1-Strep-TlaA2-Myc | pET15bTEV producing TslA lacking the first 276 amino acids as an N-terminal His(6)-fusion, alongside TlaA1 with a C-terminal strep tag and TlaA2 with a C-terminal Myc tag | This work |
| pET15bTEV-His-TilA-TslA-TlaA1-Strep-TlaA2-Myc | pET15bTEV producing TilA lacking the first 39 amino acids as an N-terminal His(6)-tag fusion, alongside full length TslA, TlaA1 with a C-terminal strep tag and TlaA2 with a C-terminal Myc tag | This work |
| pET15bTEV-His-TilA | pET15bTEV producing TilA lacking the first 40 amino acids as an N-terminal His(6)-tag fusion | This Work |
| pET15bTEV-His-TslA$_{NT(1-311)}$ | pET15bTEV producing the N-terminal domain of TslA (aa 1 – 311) with an N-terminal His(6)-tag | This Work |
| pUT18 | Vector encoding T18 fragment of *B. pertussis* CyaA; amp$^{r}$ | Reference[14] |
| pUT18-NarG | aa 1-42 of *E. coli* NarG fused to the N-terminus of T18 CyaA | Reference[15] |
| pUT18-TilA | Mature region of TilA (i.e. lacking aa 1-24) fused to the N-terminus of T18 CyaA | This Work |
| pUT18-TslA | TslA fused to the N-terminus of T18 CyaA | This Work |
| pUT18-TslA$_{NT}$ | N-terminal domain of TslA (aa 1-260) fused to the N-terminus of T18 CyaA | This Work |
| pUT18-TslA$_{CT}$ | C-terminal domain of TslA (aa 239-442) fused to the N-terminus of T18 CyaA | This Work |
| pUT18-TlaA1 | TlaA1 fused to the N-terminus of T18 CyaA | This Work |
| pUT18-TlaA2 | TlaA2 fused to the N-terminus of T18 CyaA | This Work |
| pT25 | Vector encoding T25 fragment of *B. pertussis* CyaA cml$^{r}$ | Reference[16] |
| pT25-NarJ | *E. coli* NarJ fused to the C-terminus of T25 CyaA | Reference[15] |
| pT25-TilA | Mature region of TilA (i.e. lacking aa 1-25) fused to the C-terminus of T25 CyaA | This Work |
| pT25-TslA | TslA fused to the C-terminus of T25 CyaA | This Work |
| pT25-TslA$_{NT}$ | N-terminal domain of TslA (aa 2-260) fused to the C-terminus of T25 CyaA | This Work |
| pT25-TslA$_{CT}$ | C-terminal domain of TslA (aa 239-442) fused to the C-terminus of T25 CyaA | This Work |
| pT25-TlaA1 | TlaA1 fused to the C-terminus of T25 CyaA | This Work |
| pT25-TlaA2 | TlaA2 fused to the C-terminus of T25 CyaA | This Work |

**Supplementary Table 3.** Plasmids used in this work.

| Primer | Sequence (5'-3') | Usage |
|---|---|---|
| C164 | AAACGGATTAGAATTCCTGCAGCCCGGG | Construction of pIMAY-Z-tslA |
| C165 | CATGCCTTCAGATATCAAGCTTATCGATACCGTCGAC | |
| C166 | GCTTGATATCTGAAGGCATGGTGTTATATC | |
| C167 | TCTCACCTTCAGACGGTCCAAAATTCAAATAATCT | |
| C168 | ATTTTGGACCGTCTGAAGGTGAGATGTTTAATCAAATTAATAATAAAAA TGAATTAGA | |
| C169 | GCAGGAATTCTAATCCGTTTTCAATATCTTTG | |
| C368 | TGTTGAGTAGGAATTCGTAATCATGTCATAGC | Construction of pTH100-tilA |
| C369 | AATATTTCATAAATAATCATCCTCCTAAGG | |
| C370 | GGATGATTATTTATGAAATATTCAAATGTTTATAAAATTAAAAC | |
| C371 | TTACGAATTCCTACTCAACATTCTCACC | |
| FOR_406_pRAB11_KnpI | GCGCGGTACCAGGAGGTTTCTAGTTATGTTGAGTAGGAAGTATAAAAT AG | Construction of pRAB11-TslA |
| REV_406_pRAB11_SacI | GCGCGAGCTCTCATAGTAATCCACCTATTTGTGATGC | |
| C93 | TAATAGCTAAGAGCTCGAATTCACTGGC | Construction of pRAB11-TslA-TlaA1-TlaA2 |
| C94 | GAAACCTCCTGGTACCATCATACTCTATCAATG | |
| C95 | TGATGGTACCAGGAGGTTTCTAGTTATGTTG | |
| C96 | ATTCGAGCTCTTAGCTATTAATTTTTTTCAGGTC | |
| S164A fwd | GCACTAGGTGGAAGAGATGC | TslA S164A substitution |
| S164A rev | ATGTCCAGTAATAAAATCAATGTCATACTTAC | |
| D224A fwd | GCTGCATTGACAGAAAATCTG | TslA D224A substitution |
| D224A rev | ATTTTCTGCAACAAACCTAGTAATATTACC | |
| H251A fwd | GCTGAAATGGAAGGCTTTCTG | TslA H251A substitution |
| H251A rev | ACCTTTACCATTTTTAAAGACTTTATC | |
| pRAB11_1 | TTTGCAATAAGAATTCACTGGCCGTCGTTTTAC | Construction of pRAB11-pep86-EsxA |
| pRAB11_2 | GCCGGAGCCGCTAATTTTTTTAAACAGGCGCCAGCCGCTCACCATAACT AGAAACCTCCTGGTACCATCATACTCTATCAATGATAG | |
| esxA_fwd | AGGAGGTTTCTAGTTATGGTGAGCGGCTGGCGCCTGTTTAAAAAAATT AGCGGCTCCGGCGCAATGATTAAGATGAGTCC | |
| esxA_rev | CAGTGAATTCTTATTGCAAACCGAAATTATTAG | |
| C227 | CGCCTGTTTAAAAAAATTAGCGGCTCCGGCTTGAGTAGGAAGTATAAA ATAG | Construction of pRAB11-pep86-TslA-TlaA1-TlaA2 |
| C228 | CTAATTTTTTTAAACAGGCGCCAGCCGCTCACCATAACTAGAAACCTCCT GGTACC | |
| C285 | AATTTTTTTAAACAGGCGCCAGCCGCTCACCATTTAATCTCCTCC | Construction of pRAB11-TslA-TlaA1-pep86-TlaA2 |
| C286 | CGCCTGTTTAAAAAAATTAGCGGCTCCGGCACTCTAATAGAACCAG | |
| pQE70_406_SphI_fwd | CGCGGCATGCTGAGTAGGAAGTATAAAATAG | Construction of pQE70-TslA-His |
| pQE70_406_BamHI_rev | CGCGGGATCCTAGTAATCCACCTATTTGTG | |
| NM 6 | GCTAGCGCCGCCCTGAAAATACAGGT | Construction of pET15bTEV-His-TslA-TlaA1-Strep-TlaA2-Myc |
| NM 7 | GCTGAGCAATAACTAGCATAAC | |
| NM 24 | TGCTAGTTATTGCTCAGCTTACAG | |
| NM 27 | TATTTTCAGGGCGGCGCTAGCATGTTGAGTAGGAAGTATAAAATAG | |
| NM 40 | GCCACCCGCAGTTCGAAAAGTGAGCATGATTTGTTTAACTTTAAG | |
| NM 41 | TTTCGAACTGCGGGTGGCTCCAGTCACTTAGCTTATCTAACTG | |
| C333 | GAAGAAAATAATAAGTCATTTGTAAAGAATTCAAATAATGCTATCTC | Construction of pET15bTEV-His- |
| C334 | CATGCTAGCGCCGCCCTGAAAATAC | |

| | | |
|---|---|---|
| | | TslA$_{CT}$-TlaA1-Strep-TlaA2-Myc |
| NM 17 | CATGCTAGCGCCGCCCTGAAAATAC | Construction of pET15bTEV-His-TilA-TslA-TlaA1-HA-TlaA2-Myc |
| NM 18 | AGCGTAATCTGGTACGTCGTATGGGTAGTCACTTAGCTTATCTAACT | |
| NM 19 | GTTTAACTTTAAGAAGGAGACCCGGGATGATGTTTAATCAAATTAATAATA | |
| NM 20 | AGCGTAATCTGGTACGTCGTATGGGTAGTCACTTAGCTTATCTAACT | |
| NM 6 | TACGACGTACCAGATTACGCTTGAGCATGATTTGTTTAACTTTAAG | Construction of pET15bTEV-His-TilA-TslA |
| NM 7 | TTCAGAAATAAGTTTTTGTTCCATATGGCTATTAATTTTTTTCAG | |
| NM 8 | TATTTTCAGGGCGGCGCTAGCATGACAGATTCAAAAGAAGAACAAA | |
| NM 9 | CCTTCTTAAAGTTAAACAAAACCGGTACCGGTCTACTCAACATTCTCACCTTCA | |
| NM 10 | TTTGTTTAACTTTAAGAAGGAGAGGATCCATGTTGAGTAGGAAGTATAAAATAG | |
| NM 11 | GGGTTATGCTAGTTATTGCTCAGCTCATAGTAATCCACCTATTTGTG | |
| NM 13 | AAAACTTATTTCTGAAGAAGACCTGTAAGCTGAGCAATAACTAGCAT | Construction of pET15bTEV-His-TilA-TslA-TlaA2-Myc |
| NM 14 | GCTGAGCAATAACTAGCATAAC | |
| NM 15 | TTTGTTTAACTTTAAGAAGGAGACCCGGGATGACTCTAATAGAACCAGAT | |
| NM 16 | GTCTTCTTCAGAAATAAGTTTTTGTTCCATATGGCTATTAATTTTTTTCAGGTC | |
| NM 6 | GCTAGCGCCGCCCTGAAAATACAGGT | Construction of pET15bTEV-His-TilA |
| NM 7 | GCTGAGCAATAACTAGCATAAC | |
| NM 8 | TATTTTCAGGGCGGCGCTAGCATGACAGATTCAAAAGAAGAACAAA | |
| NM 85 | GCTAGTTATTGCTCAGCCTACTCAACATTCTCACCTTCA | |
| NM 6 | GCTAGCGCCGCCCTGAAAATACAGGT | pET15bTEV-His-TslA$_{NT(1-311)}$ |
| NM 7 | GCTGAGCAATAACTAGCATAAC | |
| NM 27 | TATTTTCAGGGCGGCGCTAGCATGTTGAGTAGGAAGTATAAAATAG | |
| NM 95 | ATGCTAGTTATTGCTCAGCTCATCCTCCATTTGTAGTCATCATA | |
| FOR_405_pT18_BamHI | GCGCGGATCCCATGGAAATGATGGAGTAT | Construction of pUT18-TilA |
| REV_405_pT18_KpnI | GCGCGGTACCCGCTCAACATTCTCACCTTCAGACGG | |
| FOR_406_pT18_BamHI | GCGCGGATCCCATGTTGAGTAGGAAGTATAAAATAG | Construction of pUT18-TslA |
| REV_406_pT18_KpnI | GCGCGGTACCCGTAGTAATCCACCTATTTGTGATGC | Construction of pUT18-TslA/pUT18-TslA$_{CT}$ |
| pUT18_406_D2 fwd | GCGCGGATCCCATG GGTAATGATAAAGTCTTTAA | Construction of pUT18-TslA$_{CT}$ |
| C158 | TTCTTCGGTCAGAAAGCC | Construction of pUT18-TslA$_{NT}$ |
| C159 | CGGGTACCGAGCTCGAATTC | |
| pUT18_407_Fwd | GCGCGGATCCCATGATGTTTAATCAAATTAATAATAAAAATG | Construction of pUT18-TlaA1 |
| pUT18_407_Rev | GCGCGGTACCCGGTCACTTAGCTTATCTAACTGC | |
| pUT18_408_Fwd | GCGCGGATCCCATGACTCTAATAGAACCAG | Construction of pUT18-TlaA2 |
| pUT18_408_Rev | GCGCGGTACCCGGCTATTAATTTTTTTCAGGTCATCC | |
| FOR_405_pT25_BamHI | GCGCGGATCCCGAAATGATGGAGTAT | Construction of pT25-TilA |
| REV_405_pT25_KpnI | GCGCGGTACCCTACTCAACATTCTCACCTTC | |

| | | |
|---|---|---|
| FOR_406_pT25_BamHI | GCGCGGATCCCTTGAGTAGGAAGTATAAAATAG | Construction of pT25-TslA |
| REV_406_pT25_KpnI | GCGCGGTACCTCATAGTAATCCACCTATTTG | Construction of pT25-TslA and pT25-TslA$_{CT}$ |
| pT25_406_D2fwd | GCGCGGATCCC GGTAATGATAAAGTCTTTAA | Construction of pT25-TslA$_{CT}$ |
| C161 | TTCTTCGGTCAGAAAGCCTTCCATTTC | Construction of pT25-TslA$_{NT}$ |
| C162 | TAGGGTACCTAAGTAAGTAAGAATTCA | |
| pT25_407_Fwd | GCGCGGATCCCATGTTTAATCAAATTAATAATAAAAATGAATTAG | Construction of pT25-TlaA1 |
| pT25_407_Rev | GCGCGGTACCTCAGTCACTTAGCTTATCTAAC | |
| pT25_408_Fwd | GCGCGGATCCCACTCTAATAGAACCAGATATG | Construction of pT25-TlaA2 |
| pT25_408_Rev | GCGCGGTACCTTAGCTATTAATTTTTTTCAGG | |

**Supplementary Table 4.** Oligonucleotides used in this work. Underlined sequences indicate restriction endonuclease sites.

| Protein | Expression time | Sample application AC | Buffer | Column |
|---|---|---|---|---|
| TslA (wildtype and point substituted variants) | 4h | 2.5 ml min$^{-1}$ supplemented with 20 mM imidazole | Buffer A: 50 mM HEPES pH 7.5 300 mM NaCl Eluted with a gradient of 0 - 500 mM imidazole in Buffer A | 5 ml HisTrap FF |
| | | | Buffer B: 20 mM HEPES pH 7.5150 mM NaCl | Superdex® 75 pg 16/600 |
| TslA$_{CT}$-TlaA1-TlaA2 | 4h | 2.5 ml min$^{-1}$ supplemented with 50 mM imidazole | Buffer A: 20 mM Tris HCl pH 7.5, 50 mM KCl, 10% glycerol (v/v) Eluted with a gradient of 50 - 550 mM imidazole in Buffer A | 5 ml HisTrap FF |
| | | | Buffer B: 50 mM HEPES pH 8, 150 mM NaCl | Superdex® 75 pg 16/600 |
| TilA-TslA-TlaA1-TlaA2 | 2.5h | 0.5 ml min$^{-1}$ supplemented with 20 mM imidazole | Buffer A: 50 mM HEPES pH 7.5, 300 mM NaCl Eluted with a gradient of 0 - 500 mM imidazole in Buffer A | 1 ml HisTrap, FF |
| | | | Buffer A: 50 mM HEPES pH 7.5, 300 mM NaCl Step elution with 5 mM *d*-desthiobiotin in Buffer A | 1ml StrepTrap, FF |
| | | | Buffer B: 20 mM HEPES pH 7.5, 150 mM NaCl, | Superdex® 200 10/300 GL |
| TilA | 2h | 2.5 ml min$^{-1}$ supplemented with 20 mM imidazole | Buffer A: 50 mM HEPES pH 7.5, 300 mM NaCl Eluted with a gradient of 0 - 500 mM imidazole in Buffer A | 5 ml HisTrap, FF |
| | | | Buffer B: 20 mM HEPES pH 7.5, 150 mM NaCl | Superdex® 75 pg 10/300 |
| 11S-His | 4h | 2.5 ml min$^{-1}$ supplemented with 50 mM imidazole | Buffer A: 20 mM Tris HCl pH 7.5, 50 mM KCl, 10% glycerol (v/v) Eluted with a gradient of 50 - 550 mM imidazole in Buffer A | 5 ml HisTrap FF |
| | | | Buffer B: 50 mM HEPES pH 8, 150 mM NaCl | Superdex® 75 pg 16/600 |

**Supplementary Table 5.** Protein expression and purification conditions used in this work.

### Supplementary Figure Legends

**Supplementary Fig. 1. SAOUHSC_00406 has a polymorphic N-terminal region.** SAOUHSC_00406 homologues were identified by a blast search against the RefSeq database. The first 113 sequences were aligned using Muscle and visualised using Boxshade, with 13 representative sequences used to visualise the alignment of amino acids 1-296 of SAOUHSC_00406.

**Supplementary Fig. 2. Two further paralogues of SAOUHSC_00406/TslA can be encoded in *S. aureus* genomes.** a. Alignment of SAOUHSC_00406/TslA with pseduogenous paralogue, SAOUHSC_02786/TslB. b. Alignment of SAOUHSC_00406/TslA with SAPIG_RS13365/TslB and CO08_0212/TslC. Residues indicated in red were mutated to alanine as part of this study. The G-X-S-X-G motif found in some type VI-secreted lipases is indicated by the red box.

**Supplementary Fig. 3. The split nanoluciferase assay indicates that TslA is detected in the culture supernatant when co-produced with TlaA1 and TlaA2.** Whole cell samples from the same experiment as Fig. 1e were processed into a. supernatant and b. cytoplasmic fractions as described in the methods. 11S fragment of nanoluciferase and furimazine were added, and luminescence readings taken over a 10 min time course, with peak readings used to visualise the data. Experiments were performed in triplicate. Error bars are $\pm$ SD. Two-way ANOVA was used to determine statistical significance (n.s. $p>0.05$; **** $p<0.0001$).

**Supplementary Fig. 4. Bacterial two-hybrid analysis of the four proteins encoded at the *SAOUHSC_00406/tslA* cluster.** SAOUHSC_00405/TilA, SAOUHSC_00406/TslA (full length and separated N- and C-terminal domains), SAOUHSC_00407/TlaA1 and

SAOUHSC_00408/TlaA2 were produced as fusions to the T18 and T25 fragments of *Bordetella pertussis* CyaA and pairwise interactions scored in the *E. coli cyaA* mutant strain BTH101. BTH101 producing these fusions was plated onto MacConkey agar supplemented with 1% maltose, and plates were photographed after 40 hours at 30$^{o}$C. NarG-T18 and T25-NarJ were used as a positive control[15], and unfused T18 and T25 as a negative control. This experiment was repeated at least three times. Representative images are shown.

**Supplementary Fig. 5. TslA and its isolated N-terminal domain have lipase activity against the model lipid substrate Tween 20.** a. Size exclusion chromatogram of TslA-containing fractions that had been previously purified by Ni-affinity chromatography. b. SDS PAGE analysis of the peak fractions from a. The indicated fractions were pooled and used for activity assays. c. Size exclusion chromatogram of $TslA_{NT}$-containing fractions that had been previously purified by Ni-affinity chromatography. d. SDS PAGE analysis of the two peak fractions from c. Fractions from peak b were pooled and used for activity assays. e-g. Tween 20 activity assays (carried out as described in the methods) with TslA, TslA amino acid substituted variants or $TslA_{NT}$ as indicated. Experiments were performed in technical and biological triplicates. Error bars are $\pm$ SD. h. Circular dichroism spectra for purified TslA and the S164A, D224A and H251A variants.

**Supplementary Fig. 6. *S. aureus* strains encode three DUF576 tandem lipoprotein (Til1) islands and one orphan Til1.** Til1 proteins are encoded at four loci in *S. aureus* genomes[17]. The loci shown here are from the NCTC8325 genome. The hatched shading indicates a probable pseudogene. Gene diagrams were visualised in Clinker[18].

**Supplementary Fig. 7. Purification of the TilA immunity protein.** Size exclusion chromatogram of TilA-containing fractions that had been previously purified by Ni-affinity chromatography (left) and SDS PAGE analysis of the peak fraction (right).

**Supplementary Fig. 8. TslA and catalytically inactive variants are produced at similar levels in USA300 strains.** USA300 and the isogenic *til1* mutant carrying pRAB11 (vector) or pRAB11 encoding TlaA1 and TlaA2 alongside the indicated variant of TslA were grown for 2 h post induction in TSB supplemented with 5 mM $CaCl_2$. The equivalent of 1 ml of culture of $OD_{600}$ = 1 was withdrawn from each, pelleted and resuspended in PBS containing 1 mg $ml^{-1}$ lysostaphin. Samples were incubated at 37°C for 30 min and boiled for 10 min in 2 X Laemmli buffer before analysis by SDS PAGE and Western blotting with anti-TslA antibodies.

**Supplementary Fig. 9. Analysis of *S. aureus* membrane lipids following intoxication by TslA.** USA300 and USA300 Δ*til1* harbouring pRAB11 encoding TslA-TlaA1-TlaA2, were cultured for 2 h after which 500 ng $ml^{-1}$ ATc was added to induce plasmid-encoded gene expression. Samples were subsequently withdrawn at 2 and 6 h post induction and membranes prepared as described in methods. Mass spectrometric analysis, in negative ion mode (100-1000 m/z) was carried out on membranes for a. USA300 pRAB11 and b. USA300 TslA-TlaA1-TlaA2 after 2 h post induction, c. USA300 Δ*til1* TslA-TlaA1-TlaA2 after 2 h post induction and d. USA300 Δ*til1* TslA-TlaA1-TlaA2 after 6 h post induction.

```
SAOUHSC_00406/1-442     1 ----MLSRKYKIDLKAINQNTESTSAISKASYEVENANNNGLSKRDVINQFNDLKKMKKF
WP_110228616.1/1-441    1 -----MKDKYKIDPGIIKNNTEETTAISKISYEVENANLYGADSEDITRQIEYLKAKKKF
WP_001581953.1/1-441    1 -----MKDKYKIDPGIIKNNTEETTAISKISYEVENANLYGADSEDITRQIEYLKAKKKF
WP_064140200.1/1-441    1 -----MKDKYKIDPGIIKNNTEETTAISKISYEAENANLYGADSEDITRQIEYLKAKKKF
WP_165794351.1/1-428    1 ------------------NNTEETTAISKISYEVENANLYGADSEDITRQIEYLKAKKKF
WP_258414049.1/1-441    1 -----MKDKYKIDPGPIKYNTEETTAISKISYEIENAKLYGIKSKTIGRQIDQLKEAKKF
WP_078370638.1/1-442    1 ----MLSKNYKIKLNAVNSKTESTSAISKAAYEIENANNNGLTQKDLRTQINTLKNAKKF
WP_241010539.1/1-446    1 MKVKILSKNYKIKLNAVNSKTESTSAISKAAYEIENANNNGLTQKDLRTQINTLKNAKKF
WP_162665452.1/1-442    1 ----MLSSKKKIDLKAMNQNTEYTSGISKASYEVENANNNGLSKRDVINQFNDLKKMKKF
WP_248313758.1/1-442    1 ----MLSRKYKIDLKAINQNTESTSAISKASYEVENVNNNGLSKRDVINQFNDLKKMKKF
WP_084986278.1/1-442    1 ----MLSRKYKIDLKAINQNTESTSAISKASYEVENANNNGLSKRDVINQFNDLKKMKKF
WP_098691688.1/1-442    1 ----MLSRKYKIDLKAINQNTESTSAISKASYEVENANNNGLSKRDVINQFNDLKKMKKI
WP_086045600.1/1-442    1 ----MLSRKYKIDLKAINQNTESTSAISKASYEVENANNNGLSKRDEINQFNDLKKMKKF
WP_031889785.1/1-430    1 ----------------INQNTESTSAISKASYEVENANNNGLSKRDVINQFNDLKKMKKF

SAOUHSC_00406/1-442    57 PSNLEYVDSYTDSLTGVTTSAFLNKDTGKVTLGMTGTNLQDEAFKKLKEGEFSRQNVTNA
WP_110228616.1/1-441   56 PSNLEYVDSYTDSLNGVTTSAFLNKDTGKVTLGMTGTNIQGEAFKKLKEGEFSRQNVTNA
WP_001581953.1/1-441   56 PSNLEYVDSYTDSFNGVTTSAFLNKDTGKVTLGMTGTNIQGEAFKKLKEGEFSRQNVTNA
WP_064140200.1/1-441   56 PSNLEYVDSYTDSLNGVTTSAFLNKDTGKVTLGMTGTNIQGEAFKKLKEGEFSRQNVTNA
WP_165794351.1/1-428   43 PSNLEYVDSYTDSLNGVTTSAFLNKDTGKVTLGMTGTNIQGEAFKKLKEGEFSRQNVTNA
WP_258414049.1/1-441   56 PSNLEYVDSYTDSLNGVTTSAFLNKDTGKVTLGMTGTNIQGEAFKKLKEGEFSRQNVTNA
WP_078370638.1/1-442   57 PSNLEYVDSYTDSLTGVTTSAFLNKDTGKVTLGMTGTNIQGEAFKKLKEGEFSRQNVTNA
WP_241010539.1/1-446   61 PSNLEYVDSYTDSLTGVTTSAFLNKDTGKVTLGMTGTNLQDEAFKKLKEGEFSRQNVTNA
WP_162665452.1/1-442   57 PSNLEYVDSYTDSLTGVTTSAFLNKDTGKVTLGMTGTNLQDEAFKKLKEGEFSRQNVTNA
WP_248313758.1/1-442   57 PSNLEYVDSYTDSLTGVTTSAFLNKDTGKVTLGMTGTNLQDEAFKKLKEGEFSRQNVTNA
WP_084986278.1/1-442   57 PSNLECVDSYTDSLTGVTTSAFLNKDTGKVTLGMTGTNLQDEAFKKLKEGEFSRQNVTNA
WP_098691688.1/1-442   57 PSNLEYVDSYTDSLTGVTTSAFLNKDTGKVTLGMTGTNLQDEAFKKLKEGEFSRQNVTNA
WP_086045600.1/1-442   57 PSNLEYVDSYTDSLTGVTTSAFLNKDTGKVTLGMTGTNLQDEAFKKLKEGEFSRQNVTNA
WP_031889785.1/1-430   45 PSNLEYVDSYTDSLTGVTTSAFLNKDTGKVTLGMTGTNLQDEAFKKLKEGEFSRQNVTNA

SAOUHSC_00406/1-442   117 LETVKDGYADLKILYSPASDQNYRYANTQEFINKIKSKYDIDFITGHSLGGRDAVVLGMS
WP_110228616.1/1-441  116 LETVKDGYADLKILYSVLLQKKYRYANTQEFINKIKSKYDIDFITGHSLGGRDAVVLGMS
WP_001581953.1/1-441  116 LETVKDGYADLKILYSPASDQNYRYANTQEFINKIKSKYDIDFITGHSLGGRDAVVLGMS
WP_064140200.1/1-441  116 LETVKDGYADLKILYSPASDQNYRYANTQEFINKIKSKYDIDFITGHSLGGRDAVVLGMS
WP_165794351.1/1-428  103 LETVKDGYADLKILYSPASDQNYRYANTQEFINKIKSKYDIDFITGHSLGGRDAVVLGMS
WP_258414049.1/1-441  116 LETVKDGYADLKILYSPASDQNYRYANTQEFINKIKSKYDIDFITGHSLGGRDAVVLGMS
WP_078370638.1/1-442  117 LETVKDGYADIKILYSPASDQNYRYANTQEFINKIKSKYDIDFITGHSLGGRDAVVLGMS
WP_241010539.1/1-446  121 LETVKDGYADLKILYSPASDQNYRYANTQEFINKIKSKYDIDFITGHSLGGRDAVVLGMS
WP_162665452.1/1-442  117 LETVKDGYADLKILYSPASDQNYRYANTQEFINKIKSKYDIDFITGHSLGGRDAVVLGMS
WP_248313758.1/1-442  117 LETVKDGYADLKILYSPASDQNYRYANTQEFINKIKSKYDIDFITGHSLGGRDAVVLGMS
WP_084986278.1/1-442  117 LETVKDGYADLKILYSPASDQNYRYANTQEFINTIKSKYDIDFITGHSLGGRDAVVLGMS
WP_098691688.1/1-442  117 LETVKDGYADLKILYSPASDQNYRYANTQEFINKIKSKYDIDFITGHSLGGRDAVVLGMS
WP_086045600.1/1-442  117 LETVKDGYADLKILYSPASDQNYRYANTQEFINKIKSKYDIDFITGHSLGGRDAVVLGMS
WP_031889785.1/1-430  105 LETVKDGYADLKILYSPASDQNYRYANTQEFINKIKSKYDIDFITGHSLGGRDAVVLGMS

SAOUHSC_00406/1-442   177 NGIPNIVVYNPAPISITSLNPNSPDGKRLLELYKNYKGNITRFVAENDALTENLKKYKHY
WP_110228616.1/1-441  176 NGIPNIVVYNPAPISINSLNPNSPDGKRLLELYKNYKGNITRFVAENDALTENLKKYKHY
WP_001581953.1/1-441  176 NGIPNIVVYNPAPISINSLNPNSPDGKRLLELYINYKGNITRFVAENDALTENLKKYKHY
WP_064140200.1/1-441  176 NGIPNIVVYNPAPISINSLNPNSPDGKRLLELYKNYKGNITRFVAENDALTENLKKYKHY
WP_165794351.1/1-428  163 NGIPNIVVYNPAPISINSLNPNSPDGKRLLELYKNYKGNITRFVAENDALTENLKKYKHY
WP_258414049.1/1-441  176 NGIPNIVVYNPAPISINSLNPNSPDGKRLLELYKNYKGNITRFVAENDALTENLKKYKHY
WP_078370638.1/1-442  177 NGIPNIVVYNPAPISITSLNPNSPDGKRLLELYKNYKGNITRFVAENDALTENLKKYKHY
WP_241010539.1/1-446  181 NGIPNIVVYNPAPISITSLNPNSPDGKRLLELYKNYKGNITRFVAENDALTENLKKYKHY
WP_162665452.1/1-442  177 NGIPNIVVYNPAPISITSLNPNSPDGKRLLELYKNYKGNITRFVAENDALTENLKKYKHY
WP_248313758.1/1-442  177 NGIPNIVVYNPAPISITSLNPNSPDGKRLLELYKNYKGNITRFVAENDALTENLKKYKHY
WP_084986278.1/1-442  177 NGIPNIVVYNPAPISITSLNPNSPDGKRLLELYKNYKGNITRFVAENDALTENLKKYKHY
WP_098691688.1/1-442  177 NGIPNIVVYNPAPISITSLNPNSPDGKRLLELYKNYKGNITRFVAENDALTENLKKYKHY
WP_086045600.1/1-442  177 NGIPNIVVYNPAPISITSLNPNSPDGKRLLELYKNYKGNITRFVAENDALTENLKKYKHY
WP_031889785.1/1-430  165 NGIPNIVVYNPAPISITSLNPNSPDGKRLLELYKNYKGNITRFVAENDALTENLKKYKHY

SAOUHSC_00406/1-442   237 VFFGNDKVFKNGKGHEMEGFLTEEEQKAIKKELKKLQGYAEENNKSFVKNSNNAISKLAS
WP_110228616.1/1-441  236 VFFGNDKVFKNGKGHEMEGFLTEEEQKAIKKELKKLQGYAEENNKSFVKNSNNAISKLAS
WP_001581953.1/1-441  236 VFFGNDKVFKNGKGHEMEGFLTEEEQKAIKKELKKLQGYAEENNKSFVKNSNNAISKLAS
WP_064140200.1/1-441  236 VFFGNDKVFKNGKGHEMEGFLTEEEQKAIKKELKKLQGYAEENNKSFVKNSNNAISKLAS
WP_165794351.1/1-428  223 VFFGNDKVFKNGKGHEMEGFLTEEEQKAIKKELKKLQGYAEENNKSFVKNSNNAISKLAS
WP_258414049.1/1-441  236 VFFGNDKVFKNGKGHEMEGFLTEEEQKAIKKELKKLQGYAEENNKSFVKNSNNAISKLAS
WP_078370638.1/1-442  237 VFFGNDKVFKNGKGHEMEGFLTEEEQKAIKKELKKLKGYAEENNKSFVKHSNNAISKLAS
WP_241010539.1/1-446  241 VFFGNDKVFKNGKGHEMEGFLTEEEQKAIKKELKKLQGYAEENNKSFVKNSNNAISKLAS
WP_162665452.1/1-442  237 VFFGNDKVFKNGKGHEMEGFLTEEEQKAIKKELKKLQGYAEENNKSFVKNSNNAISKLAS
WP_248313758.1/1-442  237 VFFGNDKVFKNGKGHEMEGFLTEEEQKAIKKELKKLQGYAEENNKSFVKNSNNAISKLAS
WP_084986278.1/1-442  237 VFFGNDKVFKNGKGHEMEGFLTEEEQKAIKKELKKLQGYAEENNKSFVKNSNNAISKLAS
WP_098691688.1/1-442  237 VFFGNDKVFKNGKGHEMEGFLTEEEQKAIKKELKKLQGYAEENNKSFVKNSNNAISKLAS
WP_086045600.1/1-442  237 VFFGNDKVFKNGKGHEMEGFLTEEEQKAIKKELKKLQGYAEENNKSFVKNSNNAISKLAS
WP_031889785.1/1-430  225 VFFGNDKVFKNGKGHEMEGFLTEEEQKAIKKELKKLQGYAEENNKSFVKNSNNAISKLAS
```

a.

```
SAOUHSC_02786   1 MSQTEYQIKSGNIKGNSEETSTVSNISYEIENANNSGLKQNKIDKQIKKLQEKNKFPKNL
SAOUHSC_00406   1 MLSRKYKIDLKAINQNTESTSAISKASYEVENANNNGLSKRDVINQFNDLKKMKKFPSNL

SAOUHSC_02786  61 SYLKSYTDPKTGTTTSAFLNKDTGKVTLGMTGTNVHKDAILKQTFGVPSYQGYIDVSETL
SAOUHSC_00406  61 EYVDSYTDSLTGVTTSAFLNKDTGKVTLGMTGTNLQDEAFKKLKEGEFSRQNVTNALETV

SAOUHSC_02786 121 KDIGADVNIGLHSVTDKDPHYKNTQDFIKNIKKDYDIDIITGHSLGGRDAMILGMSNDIK
SAOUHSC_00406 121 KDGYADLKILYSPASDQNYRYANTQEFINKIKSKYDIDFITGHSLGGRDAVVLGMSNGIP

SAOUHSC_02786 181 HIVVYNPAPLAIKDVSGLYADQEELKKLIEKYDGHIVRFVSDEDELDAGVRNH-LYETAG
SAOUHSC_00406 181 NIVVYNPAPISITSLNPNSPDGKRLLELYKNYKGNITRFVAENDALTENLKKYKHYVFFG

SAOUHSC_02786 240 EKIVLKNGEGHAMSGILMSRTQAIILAELNKVKGYQDENNKALKSVRKQTRHRLHKVETL
SAOUHSC_00406 241 NDKVFKNGKGHEMEGFLTEEEQKAIKKELKKLQGYAEENNKSFVKNSNNAISKLASIELL

SAOUHSC_02786 300 RANWIQTTGGSLSSSQQQLLEALTALTIAEGLNQLVNEESQHLKKCIT--------RW--
SAOUHSC_00406 301 RANMMTTNGGGLSSSQQKVLESLTALTIAQSFSQLIDDEINQIKKMYNEKKKKFGKNWED

SAOUHSC_02786 350 ------------------------HIN------------------LETTGKK------RK
SAOUHSC_00406 361 AQKAGKAVGEDLSVNGVLNALDEGQVNESSMVREPEQMISAKERQLSTIGSSVSNYIMRV

SAOUHSC_02786 362 KLEMKLVKN-------------
SAOUHSC_00406 421 RLSINEIVDKDQVLASQIGGLL
```

b.

```
TslA   1 MLSRKYKIDLKAINQNTESTSAISKASYEVENANNNGLSKRDVINQFNDLKKMKKFPSNL
TslB   1 MSQTEYQIKPGNIKANSEETSTVSKISYEIENANNSGLKKGKINEQIEGLKEEGKFPKNL
TslC   1 MTKNEYKIDPGKITSNTEATSAIANISYEIENANDNGLEKEKINGQINSLKNDGDFPKNL

TslA  61 EYVDSYTDSLTGVTTSAFLNKDTGKVTLGMTGTNLQDEAFKKLKEGE-------FSRQNV
TslB  61 KYLKSYTDPNTGTTTTAFKNKDTGKVTLGMTGTNLHFEEMGNVLKHP-----FNHSDQDM
TslC  61 DYIDSYTDPSTGTTATAFLNKDTGKVTVGMAGTNFHGDQLKRVALSSMSPLLFPPSKQDM

TslA 114 TNALETVKDGYADLKILYSPASDQNYRYANTQEFINKIKSKYDIDFITGHSLGGRDAVVL
TslB 116 KDMKEMLKDFGADANIGLGAVTDKDPHFKDTQDFIKDIKKEYEIDTITGHSLGGRDAIIL
TslC 121 RDVRGTMKDGVADLAIGVGMVNYKGKHFANTQQFIENLQKKYEIDTVTGHSLGGRDAIFL

TslA 174 GMSNGIPNIVVYNPAPISITSL----------NPNSP--DGKRLLELYKNYKGNITRFVA
TslB 176 GVSNDIKNIVVYNPAPLSITDFRI----IALNNMTYSAIGDIEIKEMLKNYNGHIVRFVS
TslC 181 GLRYNIKNVVAYNPAPLEVKSIRDKFGGQLFRNTTFP--DEKYLKELMDNYDGDITKVIT

TslA 222 ENDALTENLKKYKHYVFFGNDKVFKNGKGHEMEGFLTEEEQKAIKKELKKLQGYAEENNK
TslB 232 DKDELDGFVSQFM-YDTAGEKIVLNNGEGHDMGAFLRKYAQAKILAELRKVKGYEDANNK
TslC 239 QKDGLDYLVKRTD-HLTCGDVLRINNGQGHAMENFLGEKEQREIIEELMLVKGYRDANDK

TslA 282 SFVKNSNNAISKLASIELLRANMMTTNGGGLSSSQQKVLESLTALTIAQSFSQLIDDEIN
TslB 291 SFKSVKDNTKSKLGKVEELRVNWLQANGGALTSSQLKLLESVSALIIAEGLSQLVKEESN
TslC 298 AFKALKKNTEKKLGKIDEIKSKLLQTNGGALSSSQQKLLETLVAFSVAEGLSKMVDQELQ

TslA 342 QIKKMYNEKKKKFGKNWEDAQKAGKAVGEDLSVNGVLNALDEGQVNESSMVREPEQMISA
TslB 351 QLEKMYDNMKDKFQENWKNSQEAGNKIGTKLSHDGVLTALRNGDAYESKFETEPLTKIEQ
TslC 358 QLKNMFNLMDEKFEANWKDAQEASDIVGKHLSYPEKVSALDNGGVNESKLATEPHNEIKD

TslA 402 KERQLSTIGSSVSNYIMRVRLSINEIVDKDQVLASQIGGLL
TslB 411 KLKELNDINSDCHNYVSKIKDSVNAIVGNDQLLASQIAGVI
TslC 418 KLNKITDLSKKYNSYLQQIEKSINEIVAKDQQLAGQIGDLI
```

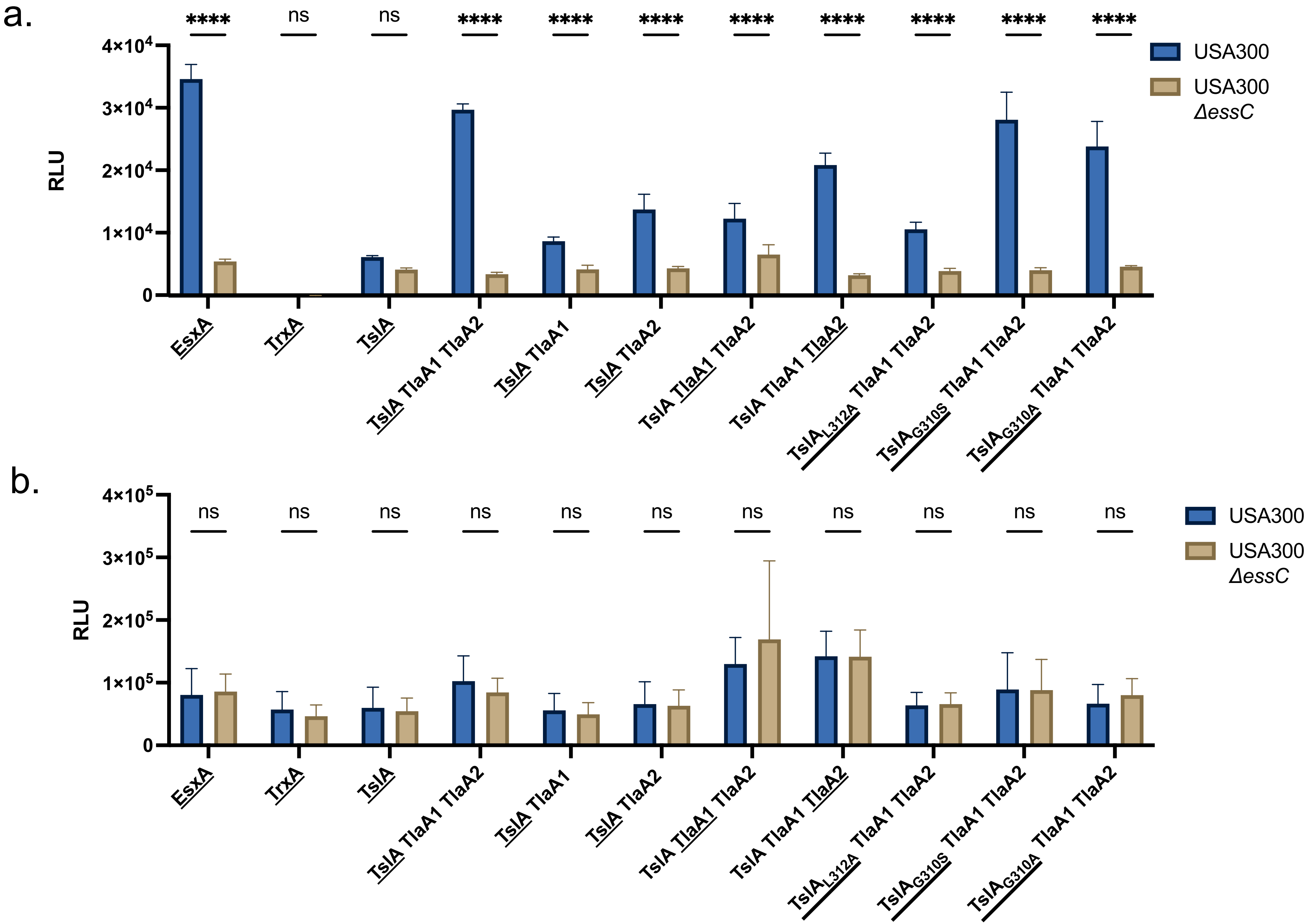

a.
b.
RLU
4×10^4
3×10^4
2×10^4
1×10^4
0
4×10^5
3×10^5
2×10^5
1×10^5
****
ns
USA300
USA300 ΔessC
EsxA
TrxA
TslA
TslA TlaA1 TlaA2
TslA TlaA1
TslA TlaA2
TslA_L312A TlaA1 TlaA2
TslA_G310S TlaA1 TlaA2
TslA_G310A TlaA1 TlaA2

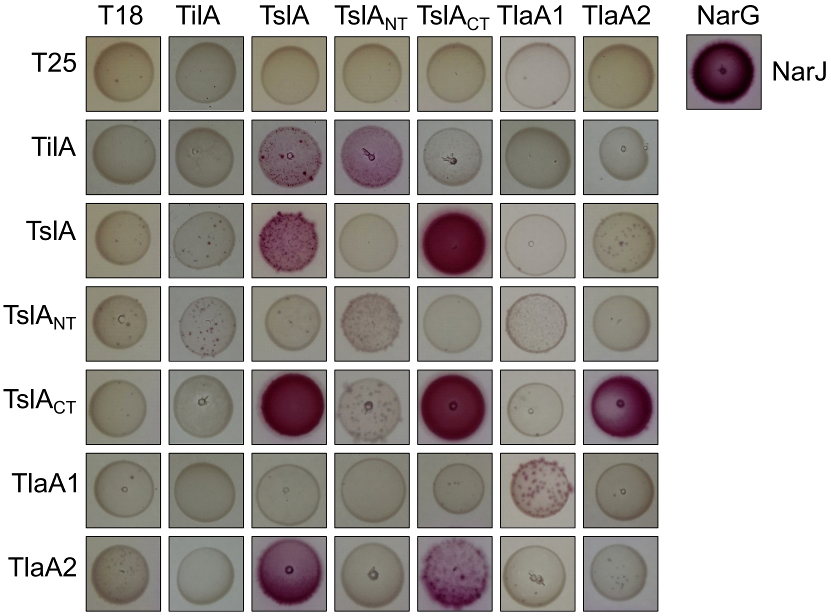
T18
TilA
TslA
TslA_NT
TslA_CT
TlaA1
TlaA2
NarG
T25
TilA
TslA
TslA_NT
TslA_CT
TlaA1
TlaA2
NarJ

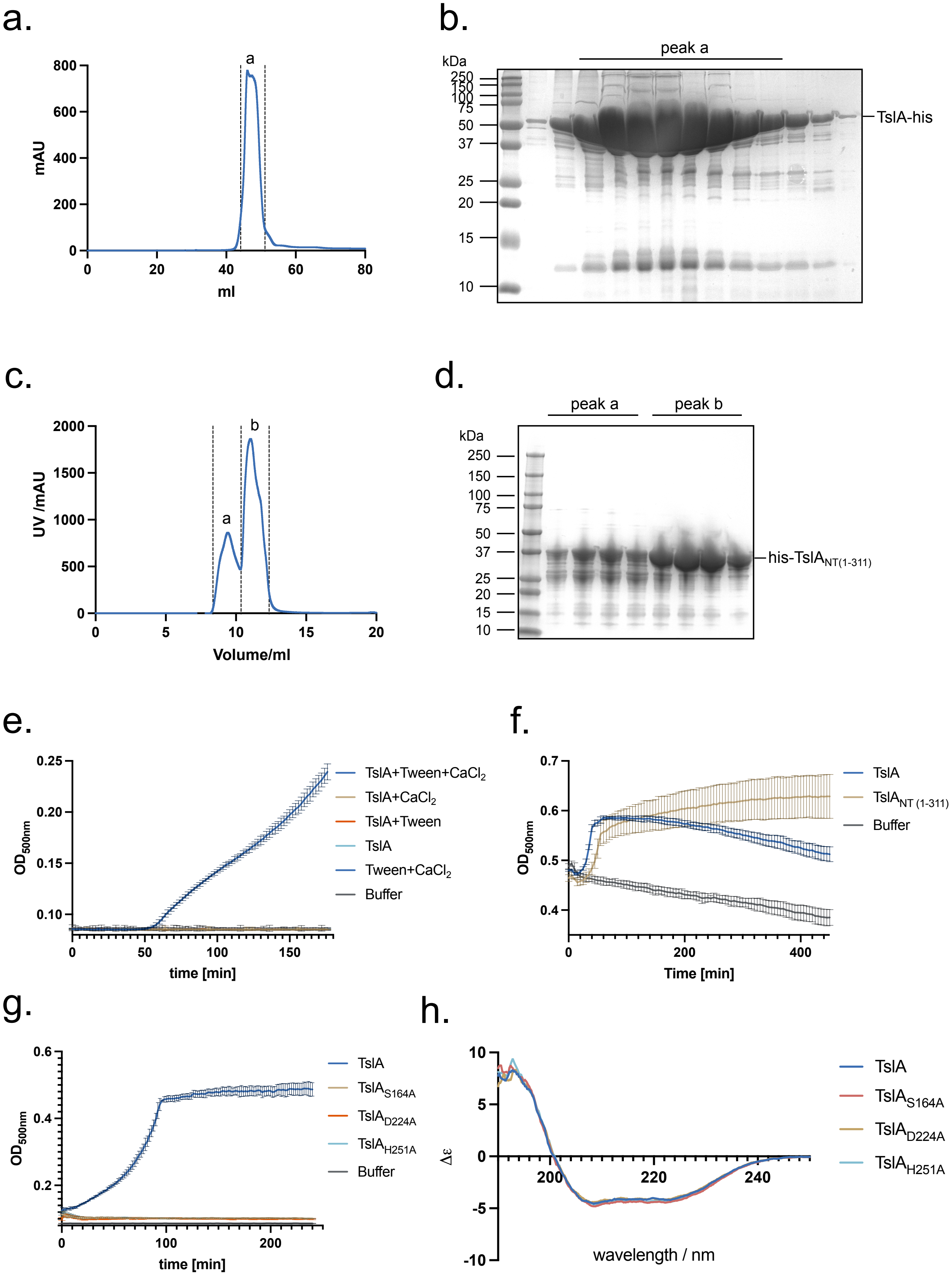

a.
a
800
600
400
200
0
mAU
0
20
40
60
80
ml
b.
peak a
kDa
250
150
100
75
50
37
25
20
15
10
TslA-his
c.
2000
1500
1000
500
0
UV /mAU
a
b
0
5
10
15
20
Volume/ml
d.
peak a
peak b
kDa
250
150
100
75
50
37
25
20
15
10
his-TslA NT(1-311)
e.
0.25
0.20
0.15
0.10
OD 500nm
0
50
100
150
time [min]
TslA+Tween+CaCl2
TslA+CaCl2
TslA+Tween
TslA
Tween+CaCl2
Buffer
f.
0.7
0.6
0.5
0.4
OD 500nm
0
200
400
Time [min]
TslA
TslA NT (1-311)
Buffer
g.
0.6
0.4
0.2
OD 500nm
0
100
200
time [min]
TslA
TslA S164A
TslA D224A
TslA H251A
Buffer
h.
10
5
0
-5
-10
Δε
200
220
240
wavelength / nm
TslA
TslA S164A
TslA D224A
TslA H251A

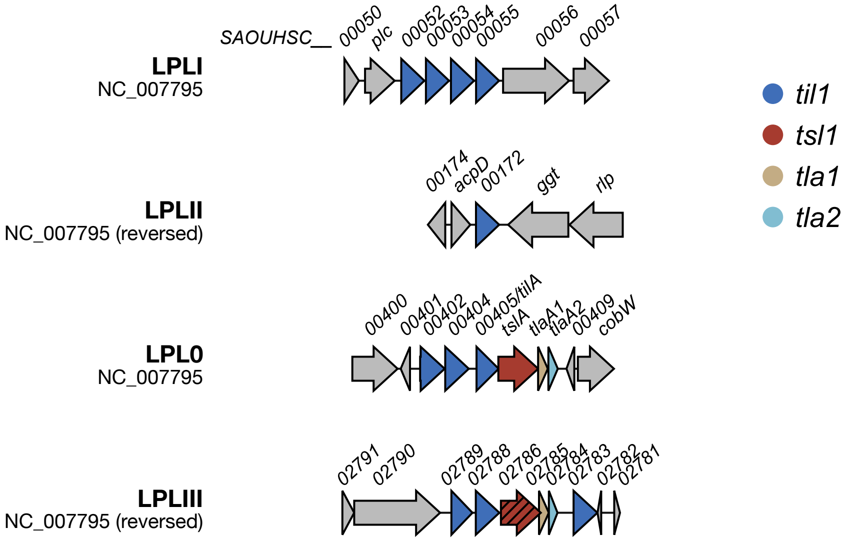
LPLI
NC_007795
SAOUHSC__
00050
plc
00052
00053
00054
00055
00056
00057
LPLII
NC_007795 (reversed)
00174
acpD
00172
ggt
rlp
LPL0
NC_007795
00400
00401
00402
00404
00405/tilA
tslA
tlaA1
tlaA2
00409
cobW
LPLIII
NC_007795 (reversed)
02791
02790
02789
02788
02786
02785
02784
02783
02782
02781
til1
tsl1
tla1
tla2

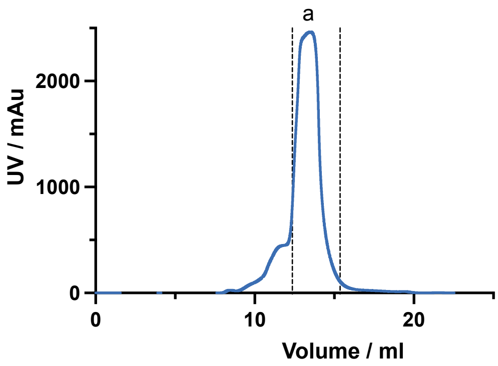
a
UV / mAu
2000
1000
0
0
10
20
Volume / ml

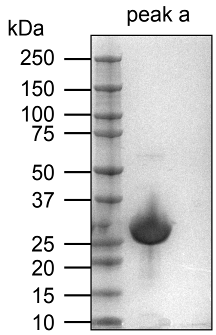
kDa
peak a
250
150
100
75
50
37
25
20
15
10

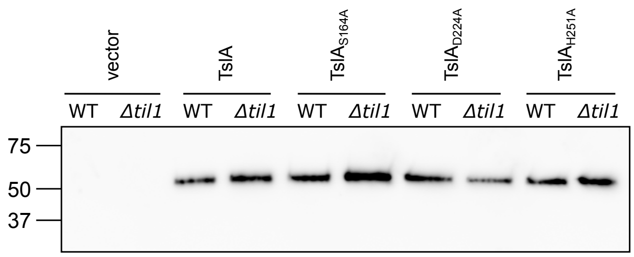

vector
TslA
$TslA_{S164A}$
$TslA_{D224A}$
$TslA_{H251A}$
WT
Δtil1
75
50
37

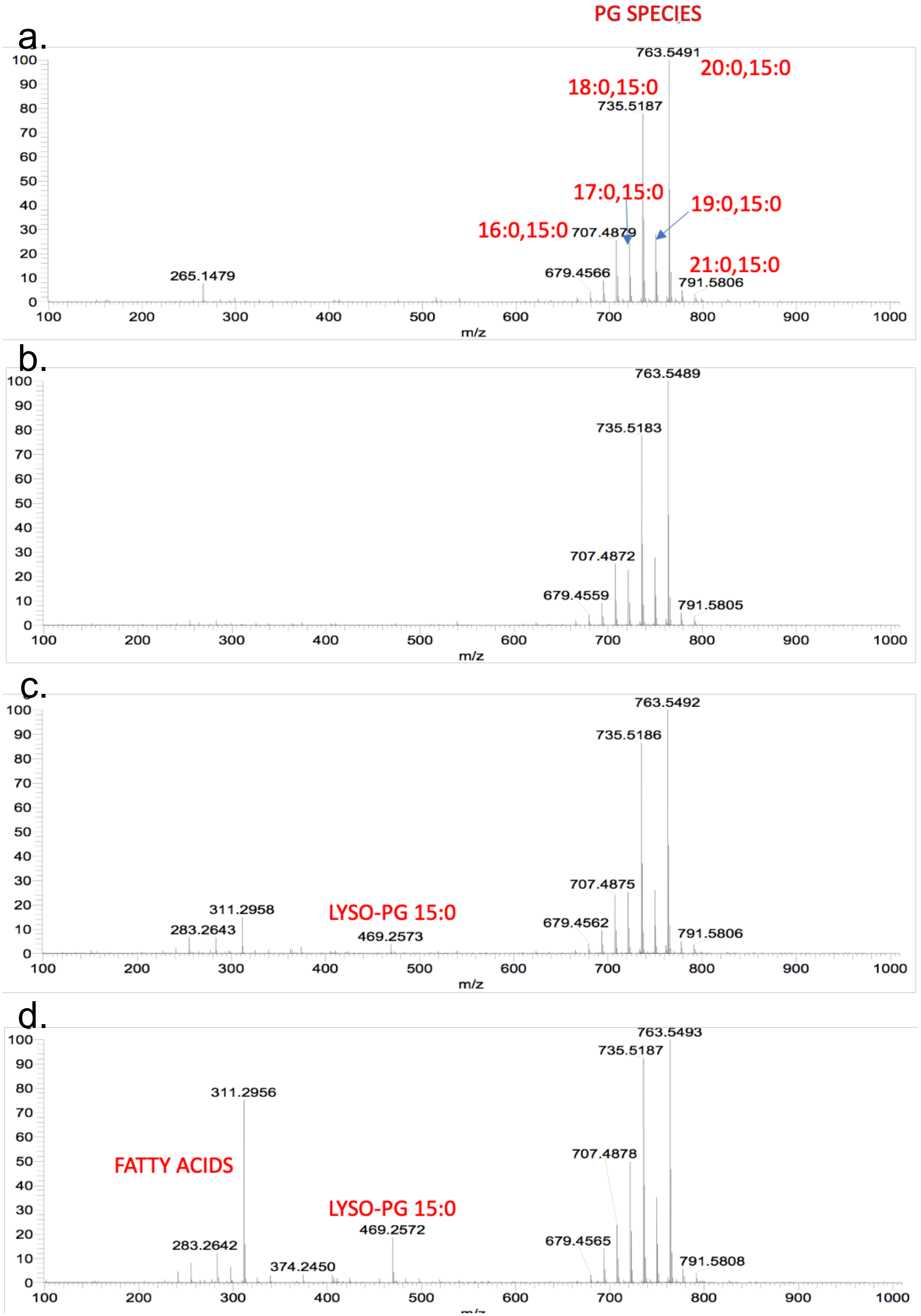
a.
b.
c.
d.
PG SPECIES
763.5491
20:0,15:0
18:0,15:0
735.5187
17:0,15:0
19:0,15:0
16:0,15:0
707.4879
21:0,15:0
679.4566
791.5806
265.1479
m/z
763.5489
735.5183
707.4872
679.4559
791.5805
763.5492
735.5186
707.4875
311.2958
LYSO-PG 15:0
283.2643
679.4562
469.2573
791.5806
763.5493
735.5187
311.2956
707.4878
FATTY ACIDS
LYSO-PG 15:0
469.2572
283.2642
679.4565
374.2450
791.5808
